## Supplementary material for "Spontaneous Chiral Symmetry Breaking in a Random Driven Chemical System": SI

### Supplementary Information

William D. Piñeros
*Center for Soft and Living Matter,*
*Institute for Basic Science (IBS), Ulsan 44919, Korea*

Tsvi Tlusty*
*Center for Soft and Living Matter,*
*Institute for Basic Science (IBS), Ulsan 44919, Korea*
*Department of Physics, Ulsan National Institute of Science and Technology (UNIST), Ulsan 44919, Korea and*
*Department of Chemistry, Ulsan National Institute of Science and Technology (UNIST), Ulsan 44919, Korea*

The following supplementary material illustrates some additional content mentioned or referenced in the main text. In particular, we provide illustrations of the case of an oscillating chemical reaction model in section I and comparable results of Figure 3 of the main text for (i) a broad range of parameters in section II and (ii) systems with inherently rugged energy landscapes in section III. We additionally discuss the distribution of individual element asymmetry of non-racemic systems in section IV, and the role of (i) noise, (ii) catalysis, and (iii) achiral to chiral element ratios and system size in sections V through VII respectively. We finally provide a detailed illustration of a chemical reaction graph for the model systems in Figure 4 of the main text in section VIII, and close with a discussion on mechanistic and interpretational aspects of the model in section IX.

*

#### I. ILLUSTRATION OF RARE OSCILLATORY SOLUTIONS

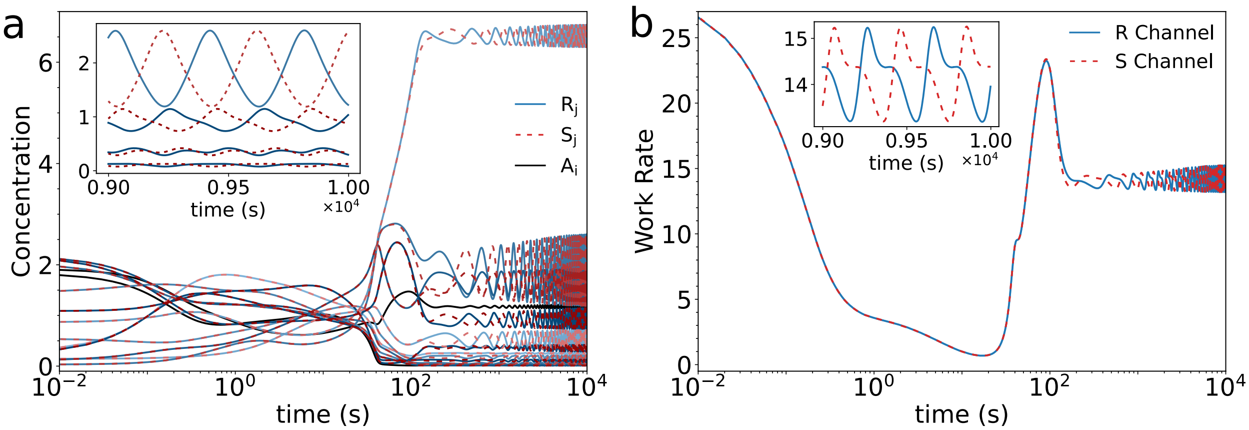


FIG. 1. An example of an oscillating system with average $ee = 0$. a. Element concentration profiles as a function of time and b. the corresponding work rate plots. Insets depict the last 1,000 steps for selected elements (a) and work rates (b) showing clear oscillatory behavior.

#### II. ILLUSTRATION OF SOLUTION DISTRIBUTIONS FOR DIFFERENT PARAMETER SETS

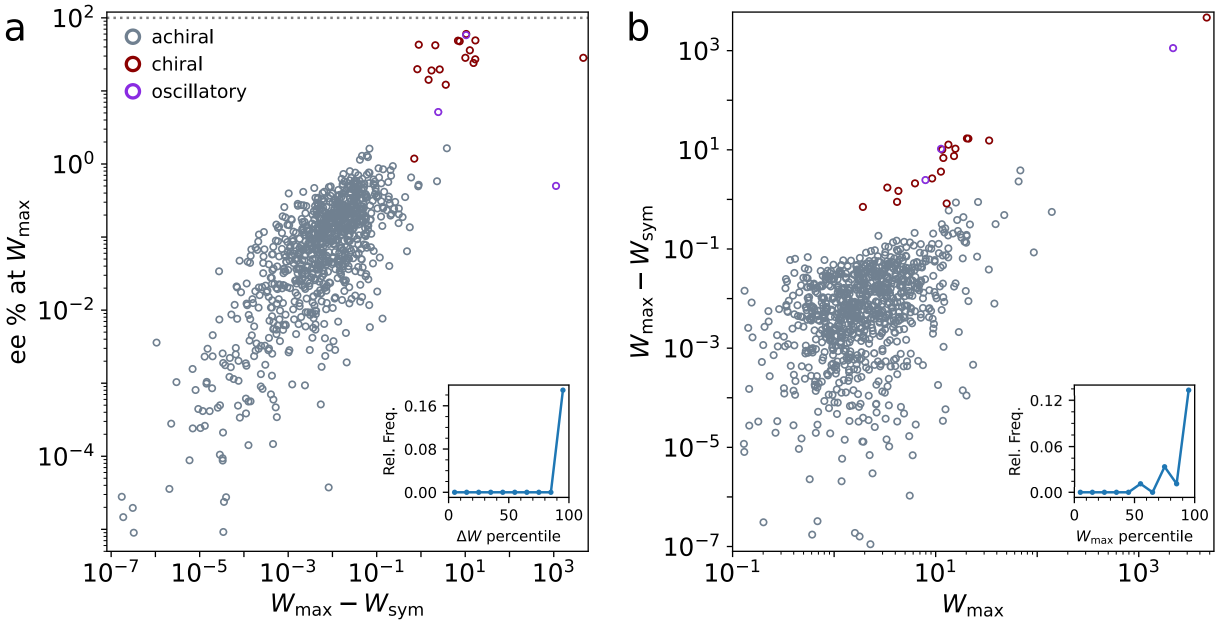


FIG. 2. a. Plot of enantiomeric excess $ee$ vs. $\Delta W = W_{\max} - W_{\text{sym}}$ and b. $\Delta W$ vs $W_{\max}$ for 900 distinct models from three different parameter sets (300 each). Points below $\sim 10^{-7}$ values cut-off for clarity. Insets: a. relative population frequency of chiral-breaking models as a function of the work difference percentile and b. the maximum work $W_{\max}$. Parameter sets are given as i) $N_a = 2$, $N_c = 2 \times 12$, $\mu = 0.3$, $s = 0.3$, $\eta = 0.3$; ii) $N_a = 8$, $N_c = 2 \times 9$, $\mu = 0.25$, $s = 0.2$,$\eta = 0.6$; and iii) $N_a = 8$, $N_c = 2 \times 9$, $\mu = 0.3$, $s = 0.3$, $\eta = 0.3$, with all sets $p_c = 0.5$, $p_b = 0.3$, $f = 10$ and $\Delta = 0.02$.

#### III. RESULTS FOR SYSTEMS WITH INHERENTLY RUGGED ENERGY LANDSCAPES

One of the original assumptions of the model presented in the main text is that of a flat equilibrium energy landscape. This was chosen to simplify implementation but is not strictly necessary. Indeed, in a more realistic and interesting scenario, the underlying energetic states are rugged and randomly distributed, especially for bimolecular reactions where unequal concentration of species might be the more likely observed case. As such, in order to incorporate a more realistic energy background, we have modified the original model to reflect random energy offsets in the underlying equilibrium reactions and repeated the analysis of the main text. In particular, reaction rate constants are now modified as

$$k_\alpha^+ = k_\alpha e^{\Delta E/2} \prod_i X_i^{a_i^\alpha} \ , \qquad k_\alpha^- = k_\alpha e^{-\Delta E/2} \prod_j X_j^{b_i^\alpha}, \tag{1}$$

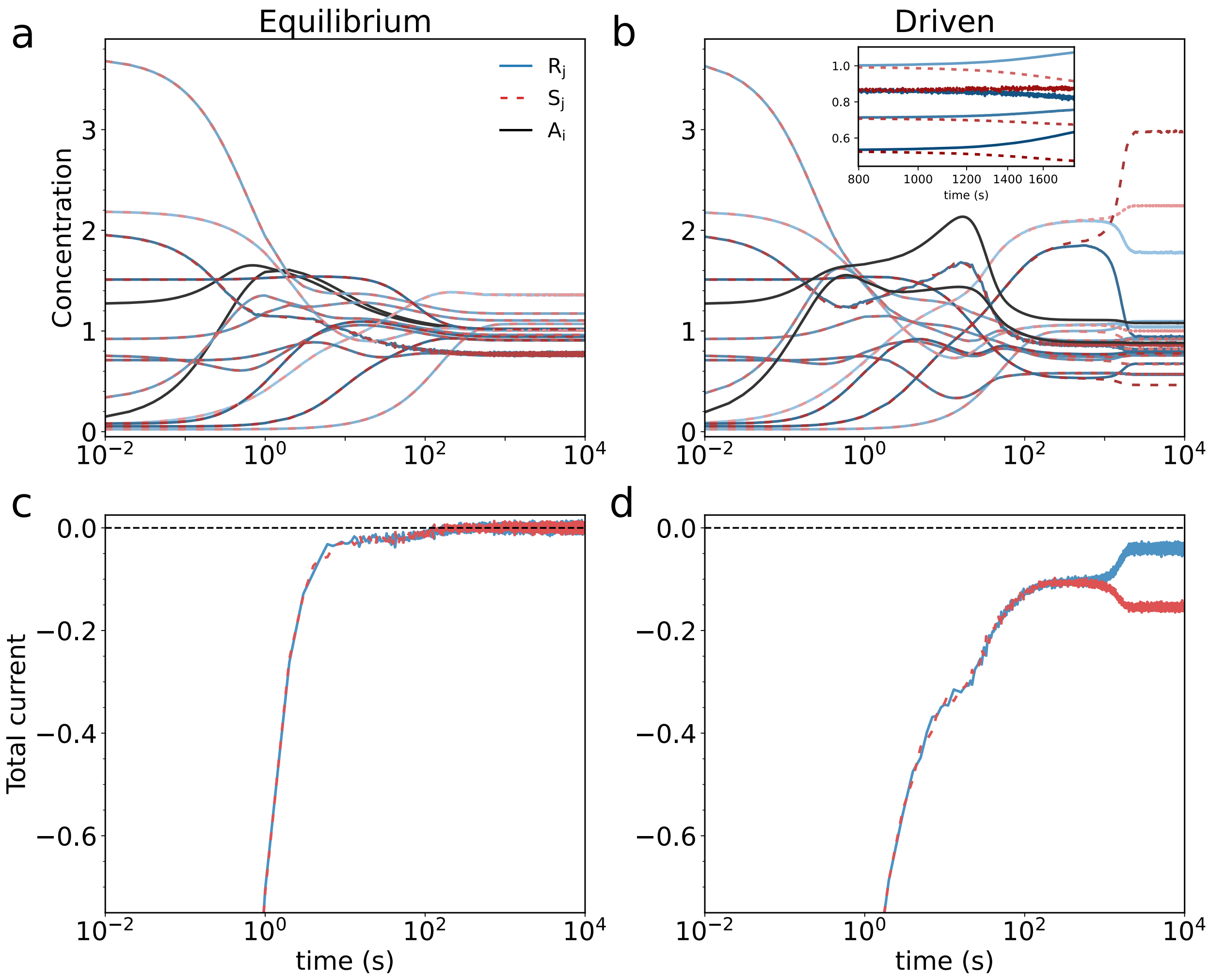


FIG. 3. a. Plot of element concentrations for a representative equilibrium system with a randomly generated, rugged energy landscape. b. Plot of the same system with identical starting conditions but subject to additional environmental forces, which in this case induce chiral symmetry breaking. c-d. Plot of total reaction current throughout the system–a proxy measure for underlying system activity–defined as $\sum_i J_i$ for all reactions.

where $\Delta E$ is chosen randomly from a uniform distribution $[-\Delta E_{\max}, \Delta E_{\max}]$ and $k_\alpha$ are determined as in the main text. Here, the energy barrier parameter $\Delta E_{\max}$ is chosen to be in the order of that of the environmental energy scale, $\sim sJ$, to maintain the system responsive to the drive *i.e.,* not trivially confined to energy wells. For the equilibrium case, not all rate parameters are free and instead must be constrained to respect detailed balance conditions as determined by the topology of the underlying network [1, 2]. Application of environmental forces, noises and simulation implementation is then carried out as per the original procedure.

Generally, we find equilibrium and driven results for this modified system replicate those of the original model, with most being moderately driven, racemic states. Yet, and as expected, we again find that these systems may indeed spontaneously break chiral symmetry

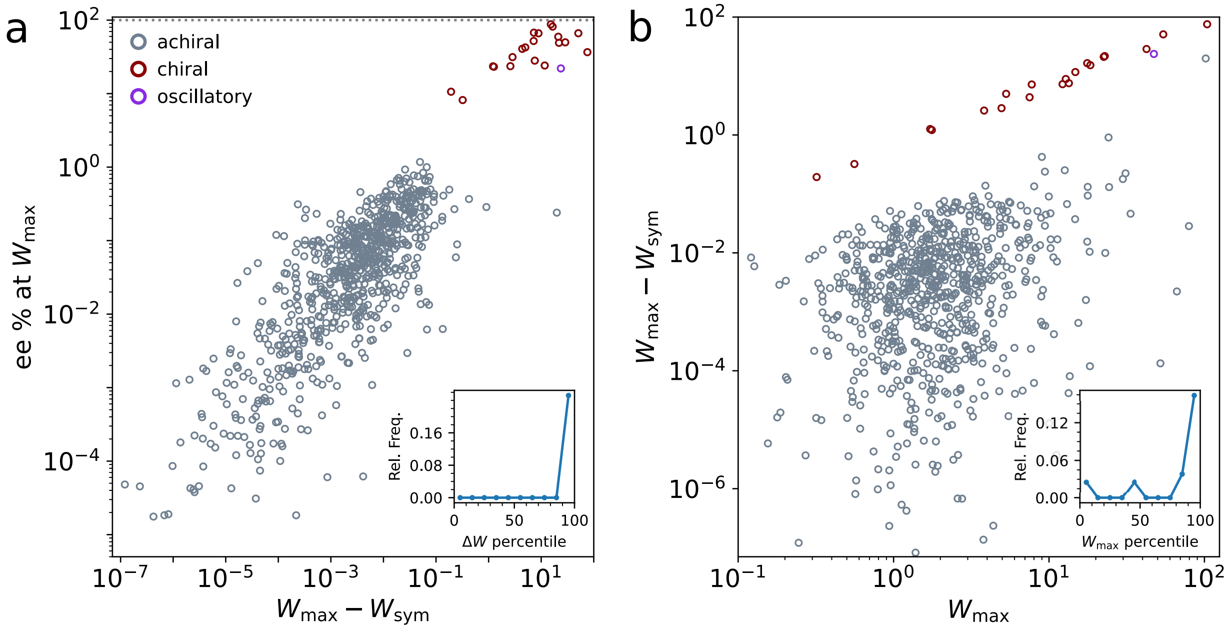


FIG. 4. a. Plot of enantiomeric excess $ee$ vs. $\Delta W = W_{\text{max}} - W_{\text{sym}}$ and b. $\Delta W$ vs $W_{\text{max}}$ for ∼800 randomly generated models following the same setting parameters as that of the main text and with the equilibrium energy landscape scale as $\Delta E_{\text{max}} = 0.2$ , *i.e.,* of approximate order as the environmental driving. Insets show the relative population frequency of chiral symmetry breaking models as a function of the work difference percentile and b. the maximum work $W_{\text{max}}$.

upon energy exploitation from an external drive. For instance, we present in figure 3a a representative equilibrium system where element concentrations are distinct due to the underlying energy landscape but nonetheless maintain detailed balance (no currents–Figure 3c). Yet, upon application of environmental forcing in this case not only drive concentrations away from equilibrium but additionally result in the spontaneous bifurcation and divergence of its chiral elements as shown in Figure 3b. As before, this is accompanied by an accompanying rise in work rate implied here through the proxy measure of net current flux in figure 3 d. to better contrast with the equilibrium case.

Carrying out a full work-harnessing analysis as before, we show in figure 4 that chiral symmetry breaking models also follow the same energy-harnessing pattern and relative frequency distribution as in figure 3 of the main text. Thus, these results show that our findings are robust to different underlying network features and energy landscapes. More importantly, they demonstrate that our original conclusions are not limited to specific assumptions but instead lend further support to a general picture of chiral symmetry breaking through energy source exploitation in driven, random chemical systems.

| | | | | | | | | | | | | |
|---|---|---|---|---|---|---|---|---|---|---|---|---|
| $R$ | 1.054 | 1.303 | 1.827 | 1.308 | 0.689 | 5.769 | 0.795 | 0.545 | 0.199 | 0.483 | 2.773 | 1.148 |
| $S$ | 0.440 | 0.645 | 0.362 | 0.244 | 0.507 | 0.172 | 1.917 | 0.628 | 0.715 | 0.537 | 0.399 | 0.323 |
| *ee* % | 41.1 | 33.8 | 66.9 | 68.5 | 15.2 | 94.2 | -41.4 | -7.1 | -56.5 | -5.3 | 74.8 | 56.1 |

TABLE I. Average steady-state concentrations and enantiomeric excess percent for the model system of Figure 2 of the main text, representing a typical system-wide asymmetry solution. The enatiomeric excess is defined as $ee = \frac{R_i - S_i}{R_i + S_i} \times 100$ for enantiomer $R_i/S_i$ pairs with negative $ee$ values denoting $S_i$ element dominance. Averages are taken over every 10th step of the last 2,000 steps of a $2.5 \times 10^4$ run.

| | | | | | | | | | | | | |
|---|---|---|---|---|---|---|---|---|---|---|---|---|
| $R$ | 0.091 | 0.083 | 0.677 | 0.606 | 0.825 | 0.812 | 0.749 | 0.088 | 1.506 | 0.071 | 0.326 | 0.856 |
| $S$ | 3.382 | 0.650 | 0.479 | 1.450 | 0.722 | 0.734 | 0.501 | 1.920 | 3.263 | 1.128 | 3.482 | 0.862 |
| *ee* % | -94.7 | -77.3 | 17.1 | -41.1 | 6.7 | 5.1 | 19.8 | -91.3 | -36.8 | -88.1 | -82.9 | **-0.34** |

TABLE II. Average steady-state concentrations and enantiomeric excess percent for a representative chiral-symmetry breaking model with near-racemic distributions in at least one of its elements ($ee < 0.5\%$, highlighted in blue). The enatiomeric excess $ee$ and average procedure are defined and calculated as in table I.

#### IV. ASYMMETRY DISTRIBUTION OF ELEMENTS IN CHIRAL-SYMMETRY BREAKING MODELS

Unlike Frank-like models with steady-state homochiral solutions, our model allows for a wide distribution of chirally-asymmetric solutions. An interesting question is what fraction of chiral elements display chiral asymmetry in a given chiral-symmetry breaking system. We generally find such systems tend to break symmetry in all elements. However, it is also possible – though rare – for some fraction of these elements to conserve chiral symmetry independent of the others. We show representative examples of each such case in table I for a typical system-wide asymmetry, and table II for a case with near-partial asymmetry where individual steady-state concentrations and corresponding $ee$ are listed.

Conceptually, these results are consistent with a picture of strong matching between system elements and environmental driving. As argued in the main text, cooperative effects take hold as a result of system-wide correlations and thus are likely to induce symmetry breaking across all chiral elements in varying proportions. On the other hand, it is also possible for some elements to be weakly associated or fully uncorrelated from the larger system, thereby conserving chiral symmetry in those elements (non-diverging dynamics). For instance consider a case where an arbitrary chiral element $R_i(S_i)$ happens to be an unreactive or inert species (*i.e.,* $\dot{R}(\dot{S})_i = 0$). Such a system will then behave as an effective $N_c - 2$ system which, by definition, can lead to chiral-breaking symmetry solutions under the right conditions of system-force matching. However, results where some elements are disconnected or weakly correlated to the rest make strong matching more difficult and hence less likely to lead to chiral-breaking dynamics. Thus such cases should be less likely to be observed while system-wide asymmetry should be more common as seen in our current results.

#### V. ROLE OF NOISE IN DYNAMICS

As briefly elaborated in the methods section, $\boldsymbol{\delta}$ represents the magnitude vector of random noises applied to unimolecular reactions of the form $A_i \rightleftharpoons R_j(S_j)$. The exact value is chosen randomly from a uniform distribution within a $[0, \Delta]$ window. Noises are therefore generically assigned but are unique for each randomly generated instance of the model. i

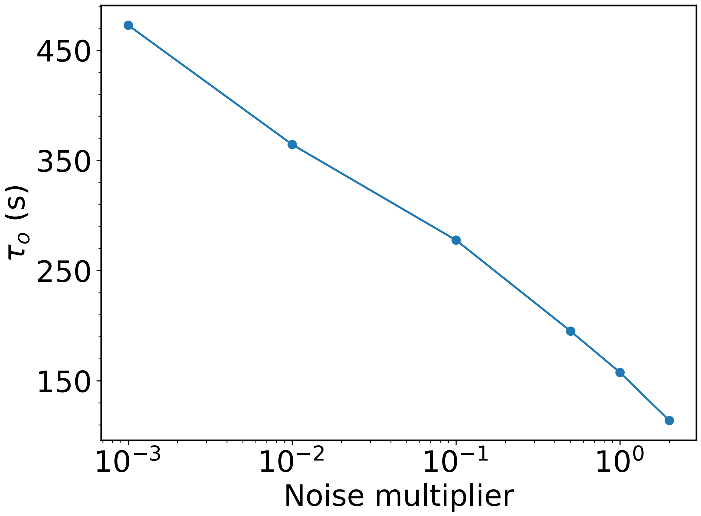


FIG. 5. Plot of bifurcation onset time $\tau_o$ vs. multiples of original system noises $\boldsymbol{\delta}$ for the same system as figure 2 in the main text. Points represent averages of 40 independent system runs with identical initial conditions. Lines are guide to the eye.

In our results, we find that most steady-state solutions lead to fixed-point solutions. As a result, noise strength is expected to only alter the onset time of bifurcation, $\tau_o$, in any given system. Indeed, as shown in Figure 5, $\tau_o$ decreases with increasing noise strength, diverging as $\boldsymbol{\delta}$ approach the noiseless limit, *i.e.,* where bifurcation cannot take place.

We can also consider how individual reaction noise alters $\tau_o$ in these systems. As discussed in the main text and section IV, given the strong correlation of elements in chiral breaking systems, we can expect that the onset time will depend largely on the magnitude of noise and not any one individual reaction. Taking as representative the same model as before, we find this system operates with $N_\delta = 6$ noisy reactions. Thus, we can systematically turn off the noise in each reaction $i$ as $\delta_i = 0$ and determine $\tau_o$ as a function of the fraction of remaining reactions. In particular, given $N_\delta$ reactions and retaining $l$ of the total leads to $N_p = \sum_{l=1}^{N_\delta - 1} \binom{N_\delta}{l}$ possible permutations of valid simulation subsets.

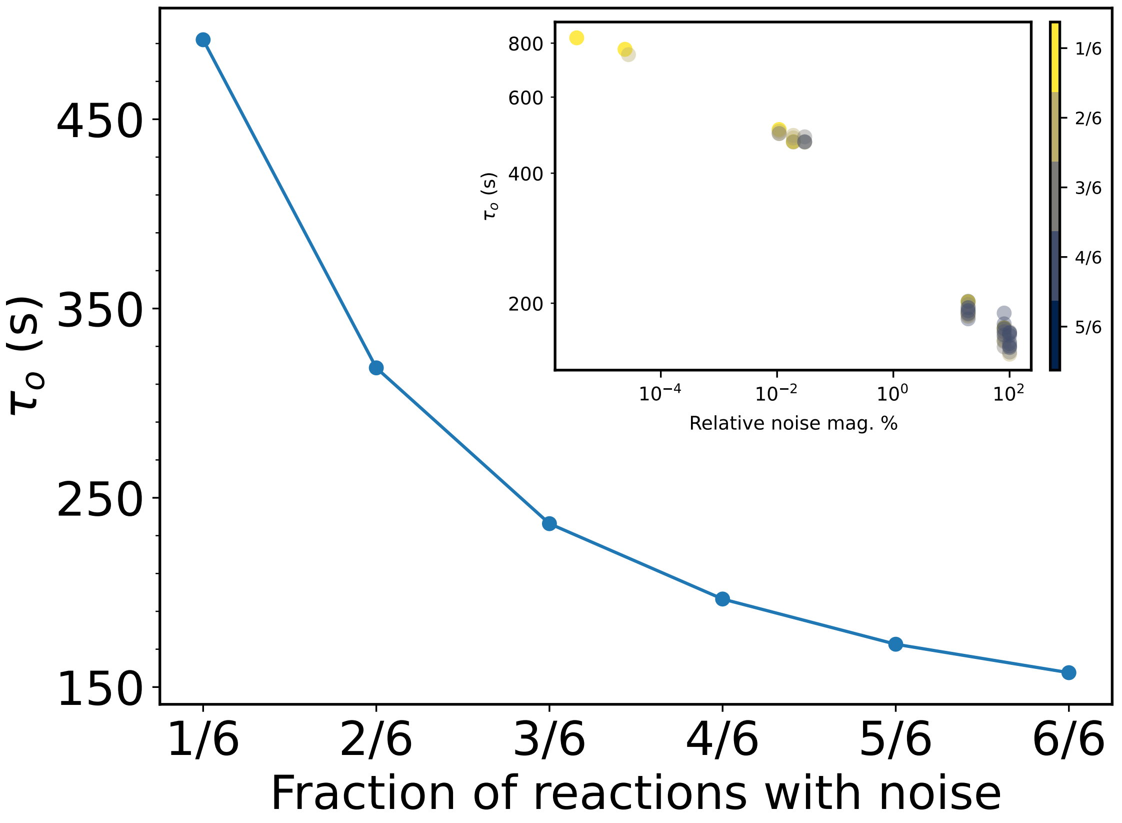


FIG. 6. Plot of bifurcation onset time vs. the number noisy reactions acting in the system. Points represent averages of $\tau_o$ from 40 independent simulations for all $\binom{N_\delta}{l}$ permutations given $l$ noisy reactions. Point for the '6/6' entry taken from the $\times 1$ multiplier entry of figure 5. Lines are guides to the eye. Inset: plot of $\tau_o$ vs. the relative magnitude percent from maximum total present for all $N_p = 62$ permutations. Color bar indicates reaction fraction membership.

We thus determine $\tau_o$ for each of the $N_p = 62$ permutations and calculate the average for each respective $l$ subset. For example, for the case where $l = 1$, 1/6 of reactions are kept and generate $\binom{6}{1} = 6$ possible cases where only $\delta_i$ of the $i = \{1, 2, ..., 6\}$ reaction is used in the simulation. A plot of these averages is shown in figure 6 showing that, as expected, $\tau_o$ increases as the fraction of noisy reactions goes down. This is understood by noting that as the fraction of noisy reactions decreases, so does the average noise felt in the system. We can show this more clearly by plotting in the figure inset $\tau_o$ against the relative noise magnitude from the maximum total for any given permutation,*i.e.*, $\sum_i^l \delta_i / \sum_i^{N_\delta} \delta_i \times 100$. Thus, as seen, $\tau_o$ varies for each permutation but depends primarily on the overall noise strength present in the system: as the noise magnitude decreases $\tau_o$ increases and roughly corresponds to a lower fraction of noisy reactions.

In all, these findings demonstrate that the overall effect of noise is to hasten bifurcation, and need not be specific to any one reaction or element. Rather the overall noise magnitude determines bifurcation time given a coordinated system response from the tightly correlated system. Similar results can be shown for other chiral symmetry-breaking models.

#### VI. ROLE OF CATALYSIS

As explained in the main text, catalysts may be added to any reaction of the system with a fixed probability $p_c$. This means that in a system of $M$ reactions, there will be, on average, $p_c M$ catalyzed reactions. In order to understand how catalysis affects the overall propensity of the model to yield chiral-symmetry breaking models, we generate and run new batches of models for different values of $p_c$ and calculate the average percentage of non-racemic models from the total. In particular, we use the original parameter set in the main text but change $p_c$ systematically as $[0, 0.25, 0.50, 0.75]$. Each modified parameter set is then used to generate new batches of 800 random systems and simulated for $2.5 \times 10^4$ steps. From this we calculate the non-racemic model percentage yield over the total number of runs. For example, a non-racemic model yield of 3% would indicate that $\sim$3 in 100 random models will be non-racemic.

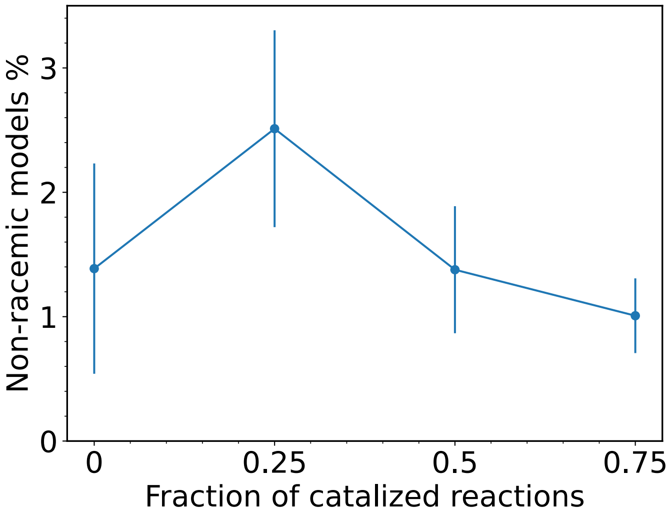


FIG. 7. Plot of average non-racemic model yield percent vs. fraction of catalyzed reactions ($p_c$ values) for the parameter set used in the main text. Each point represents average yield over a total of $\sim$800 runs. Bars indicate standard error per every 100 generated runs.

The result of this procedure is shown in figure 7. As seen, increasing the average fraction of catalyzed reactions (through $p_c$) leads to a reduction of non-racemic model yield, with a maximum around $p_c = 0.25$. More interestingly, however, chiral-symmetry breaking still takes place even without catalysts in the system. This seemingly puzzling result can be easily resolved by noting that the main chiral-symmetry breaking mechanism is that of strong matching of the forces, which introduce and maintain of non-linear feedback mechanisms in the system. In this way, catalysts may enhance the chiral-symmetry breaking process (hence the maximum) but are not strictly necessary so long as non-linear processes exist to generate and sustain the necessary divergent dynamics. Note that this result makes necessary a dynamic and configuration-dependent form of environmental driving. However, such driving can be argued to occur naturally in complex, interacting environments with steep energy forcings or gradients like those of a primordial environment.

#### VII. ACHIRAL TO CHIRAL ELEMENT DISTRIBUTION AND SYSTEM SIZE EFFECTS

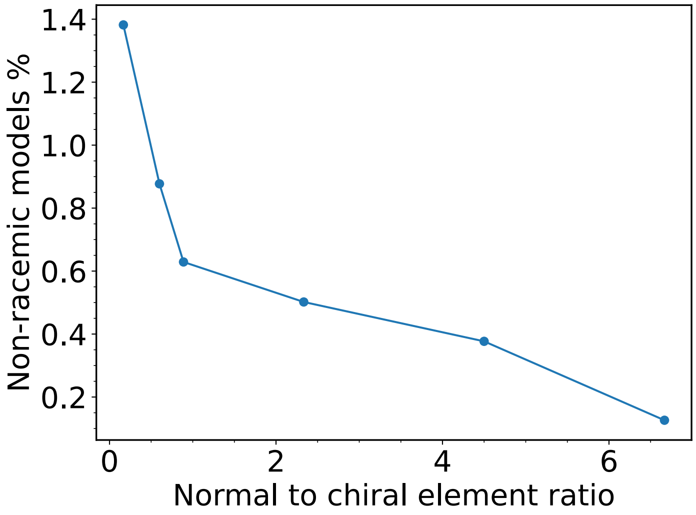


FIG. 8. Plot of average non-racemic model yield % as a function of achiral to chiral ratio $N_a : N_c$. Lines are guides to the eye.

As mentioned in the main text and previously shown in figure 2, results hold generically for a variable range of parameter values. However, to better highlight and clearly separate the effect of system size and chiral-to-achiral element ratio, we explore these conditions as systematic modifications of the original parameter set presented in the main text. As such, we first explore the achiral to chiral ratio $N_a$:$N_c$ for the original system size of $N = 26$ and determine its effect on the average non-racemic model yield percent as per section VI. In particular, we select $N_a$:$N_c$ as $[20{:}3, 18{:}4, 14{:}6, 8{:}9, 6{:}10, 1{:}12]$ where we redefine $N_c \equiv N_c/2$ to emphasize the underlying enantiomer pair symmetry (other parameters as original set). We then generate batches of 800 random models, simulate them for a total of $2.5 \times 10^4$ time steps and count the number of chiral symmetry breaking models to calculate the non-racemic yield %. Thus, as seen in figure 8, when $N_a$:$N_c$ increases, the average non-racemic model yield decreases with a sharp increase as this ratio drops below unity , *i.e.,* with excess chiral elements. These results are analogous to the findings reported in [3], where increasing the number of chiral elements in a system is more likely to result in chirally-broken steady states and attest to unstable racemic solutions.

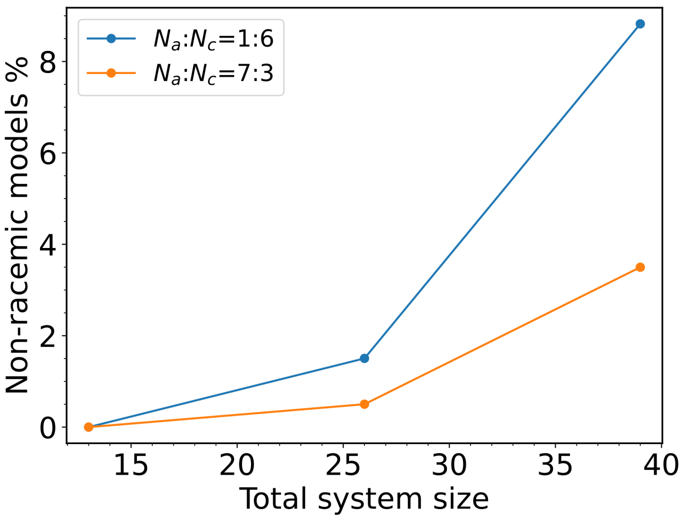


FIG. 9. Plot of average non-racemic model yield % as a function of total system size $N = N_a + 2N_c$ for a system with excess chiral elements ($N_a$:$N_c$ = 1:6, blue) and excess achiral elements ($N_a$:$N_c$ = 7:3, orange). Lines are guide to the eye.

Similarly, exploring the effect of total system size we choose $N = [13, 26, 39]$ and compare the average non-racemic model yield for a ratio with excess achiral elements $N_a$:$N_c$ = 7:3 and excess chiral elements $N_a$:$N_c$ = 1:6 (original model in main text). We then carry out the same procedure as before with 400 model batches and plot the solutions as a function of $N$ in figure 9. As seen, increasing the system size rapidly increased the average non-racemic model yield, and this effect held consistent across the two contrasting element ratios. This result is again consistent with [3] where larger system sizes, through increased chiral elements, increase the likelihood of symmetry breaking. Interestingly, no chiral symmetry breaking models were observed for the smaller systems, suggesting that a certain minimum level of complexity – attained through larger systems – is necessary to achieve the kind of system-wide correlations required in non-racemic models.

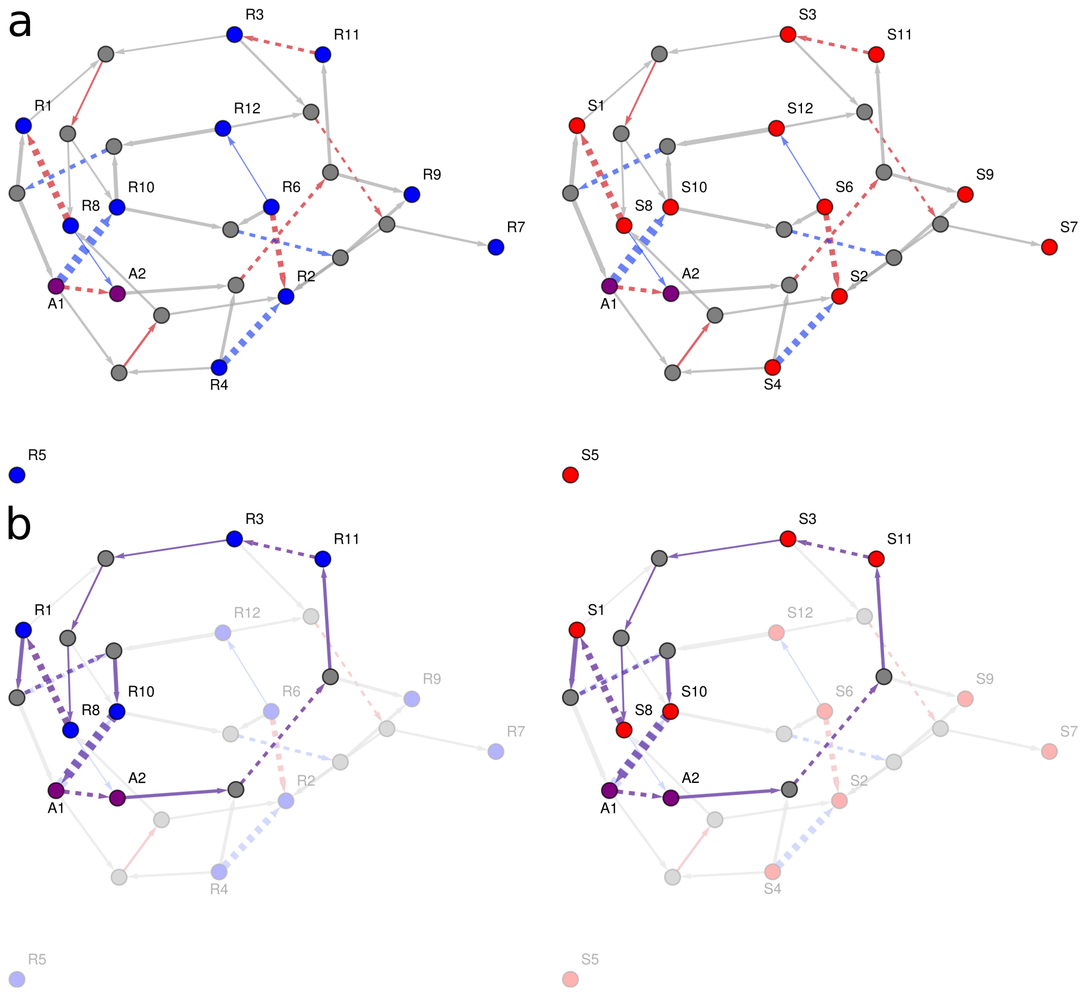


FIG. 10. a. Restricted species-reaction chemical graph for the racemic example model of figure 4 in the main text capturing 95% of currents in the system. Left represents the $R$ channel (blue elements) and right the corresponding $S$ channel (red elements). Nodes represent either elements $\{A_i, R_j, S_k\}$ (purple, blue and red respectively) or reaction complexes arising from bi-molecular reactions (gray). Colored edges represent reactions in the network with currents flowing along (red) or against (blue) the arrow direction. Edge thickness is scaled in proportion to the normalized current of the reaction $J_i' \equiv J_i/J_{\max}$ where $J_{\max}$ is the maximum magnitude value of current in the system, and $J_i$ represents the average current observed over a 2-second period during the highlighted period of rapid growth in main figure (a gray area). Gray directed edges connect elements to complexes, and thicknesses are similarly scaled to correspond in proportion to the corresponding reaction current edge. b. Highlighted cycle loop (purple) with modified edge directions to reflect current direction where applicable.

#### VIII. CHEMICAL GRAPH ILLUSTRATION OF CYCLE CURRENTS

In Figure 4 of the main text, we introduced a schematic of channel cycles to symbolize the underlying correlations between a model system during a period of rapid growth/energy harnessing. A natural question is then determining how such a network might look like and how such cycles, representing active mass currents, might manifest in the system. Here we provide one such illustration using an analogous species-complex or species-reaction chemical graph [4, 5]. These graphs differ from simple graphs as they account for intermediate nodes or "chemical complexes" that arise from the built-in chemistry of the model, such as bimolecular reactions. Thus, a reaction such as $A_i + R_j \rightleftharpoons A_k + R_l$ would introduce two new nodes or "complexes" connected by a bimolecular reaction. The complete species-complex graph is a collection of nodes representing elements and complexes, while connecting edges represent uni- and bimolecular reactions. Additionally, edges connecting elements to complexes are included to encompass the fully connected network.

For the systems in Figure 4 of the main text, these graphs result in a large number of nodes (35+) and edges (75+) so that clear illustrations, especially of cycles, are difficult and impractical. Instead, we limit the graphic analysis to those nodes and edges that carry the most current (mass flow) during the period of interest. Thus, by introducing some threshold value on the reaction currents $J_{\text{thr}}$ we reduce the complexity of the graph significantly while retaining the crucial aspects of the network.

Such graphs are shown in figures 10 and 11 for the strongly driven and chiral symmetry breaking systems of Figure4 in the main text, respectively. In particular, these graphs represent a fraction of all reactions but contain $\sim$95% and $\sim$91% of the mass flow in the system. Nonetheless, it is clear that even at this level of simplification, the reaction cycles do not manifest as a simple loop across the network but instead are tightly intertwined and allow for many possible, equivalent paths. Critically, however, tracing the current flow in these paths reveals that these tend to flow in one general direction, and that the harnessed drivings (denoted by dashed lines and thickness in proportion to the flow) act as flow pumps that power and maintain large mass flows through the entire system. For example, we highlight one such possible path for the strongly-driven racemic system in figure 10 b. As seen, 6 of these edges display forced driving along the same direction, which analogously to a circuit with low resistance, ensures large, unimpeded current flow (high energy harnessing). Further, such loops duplicate identically in the enantiomer channel so that the racemic state is maintained even in this regime.

On the other hand, looking at the chiral symmetry breaking case in figure 11a, it is obvious that the effective topology of each channel is distinct, while individually still displaying some intertwined loops powered through the environmental forces. Furthermore, unlike the racemic case where such cycles existed separately in the two channels, here the currents are system-wide and are consistent with transfer (divergence) of matter between channels. In particular, we highlight one such loop in b where the flow can be mapped to start from $A_1$ and end in $A_2$ but follow *reversed* directions within each channel. This can be seen clearly in the reaction chain from the $R_4$ to $A_2$ elements in the $R$ channel where such path not only displays stronger currents due to active driving but flows in the opposite direction in the $S$ channel. Together these graphs thus demonstrate a more detailed but consistent picture with the schematic of the main text where correlations are indicative of the strong current

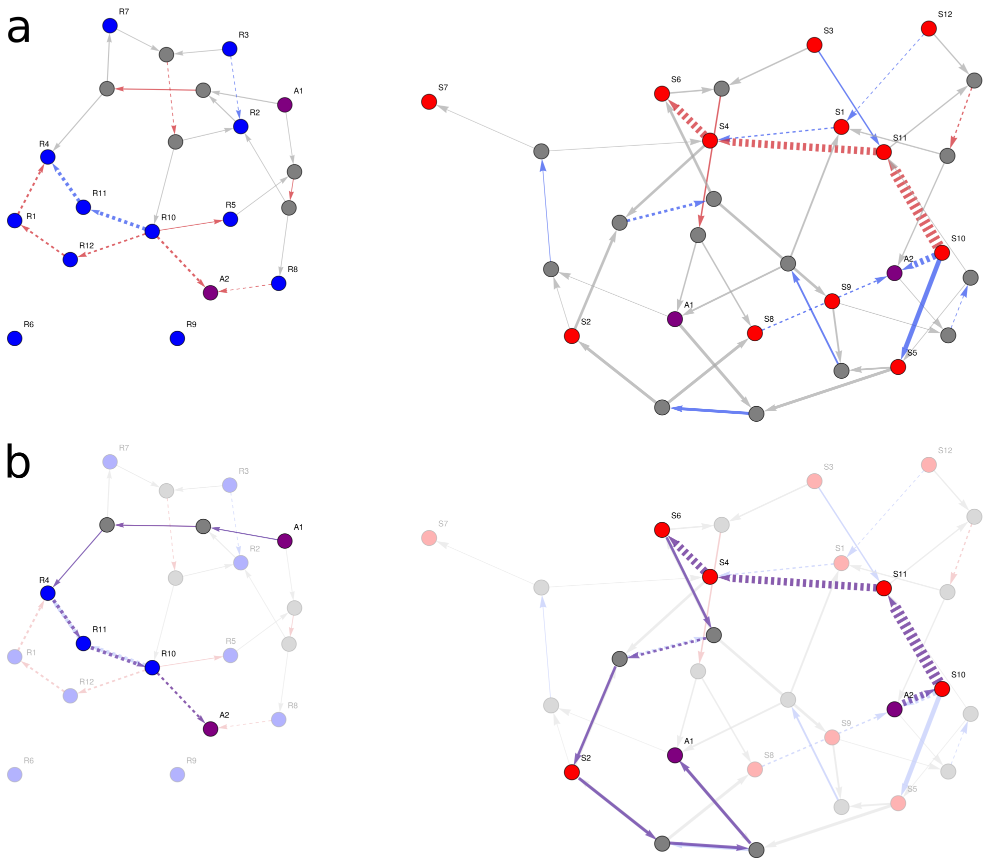


FIG. 11. a. Restricted species-reaction chemical graph for the chiral symmetry breaking model of figure 4 in the main text capturing 91% of system currents. Descriptions as in figure 10. b. Highlighted directed path showing a system-wide cycle transferring mass from the $R$ to the $S$ channel.

cycles in the network and manifest distinctly for each of the racemic and chiral symmetry breaking case.

#### IX. DISCUSSION ON THE MECHANISTIC AND INTERPRETETIONAL ASPECTS OF THE MODEL

##### A. Reaction engines: a kinetic picture

In this section we clarify mechanistic aspects of the model and discuss its interpretation within the more limited scope of reaction kinetics. In particular, as noted in the main manuscript, the model is such that a wide variety of steady-state outcomes are possible, but only a small portion of which might yield far-from-equilibrium steady states, including those leading to chiral symmetry breaking. As a simple example, consider the following minimal case from the original framework [6] involving an equilibrium reaction $A \rightleftharpoons B$ with constant forward and backward rates $k^+ = k^-$. At equilibrium inside a closed container of constant volume and temperature containing $N$ molecules, such a system will naturally settle at equal concentrations in its long-time limit. Now consider a secondary reaction pathway introduced by some feature in the environment whose *net effect* is to bias or drive reactions in some way e.g. $k'^{+} = e^{f}k^{+}$, $k'^{-} = k^{-}$ where $f(A, B)$ is the arbitrary, state dependent forcing. Then, if such driving happens to be reinforcing e.g. $f = cB$, $c > 0$, the rate of $B$ formation rapidly increases and ultimately results in a far-from-equilibrium steady-state distribution $(A, B) \sim (0, N)$. In this manner, far-from-equilibrium states may emerge from a combination of system states and driving program that enhance the rates of formation at an implicit energetic cost.

In practice, for the large, complex and randomly wired systems of this work, these far-from-equilibrium states are no longer trivial and instead manifest as strong reaction currents across dense, interconnected sets of network cycles powered by the equally random, and complex drives. Critically, even from a strict kinetic point of view, such a dramatic tilting of the element distributions is unstable and therefore requires persistent energy consumption to sustain them. Such final outcomes may then be said to describe from a chemical point of view a 'reaction engine' whose net effect is to actively 'pump' and enhance a subset of reaction elements far above equilibrium. This picture is also conceptually analogous to a recent study demonstrating distribution inversion of chemical stereomers through non-equilibrium kinetic control [7]. Given the state-dependence of the forces driving this kinetic tilting, such an 'engine' may only come about through *positive feedback* from previous configurations which thus reinforce, and maintain such driving. As a result, these type of systems *induce* their own formation—through a subsequent, implicit response to an external drive—and should be properly understood as auto-inductive.

##### B. Characterization of the feedback mechanism

Indeed, the above classification of the force mechanism is consistent with that found in chemical literature where reactions whose products enhance their own formation via some indirect, external means are termed auto-inductive [8, 9]. Such characterization may nonentheless prompt some resemblence to autocatalysis and therefore merit a closer examination in regards to our model.

In particular, general definitions of autocatalysis have been introduced in the chemical literature which define the process kinetically, through its rate equations, or stoichiometrically through the net effect along reaction channels [10, 11]. In the first, the authors define the process as $\dot{x} = kx^n + g$ where if $k, n > 0$ and $g$ small then the process is considered autocatalytic. While the authors are able to show that some elaborate examples of autocatalysis such as reaction cycles, and even some auto-induction processes, can be mapped to this form in some regimes, we note that our model excludes any such $kx^n$ terms by construction. Instead, auto-induction in our model, mapped to such an expression, comes exclusively from a complex function of $g_i$ across many elements, forcings and $i$ reactions which go beyond the original scope of that work.

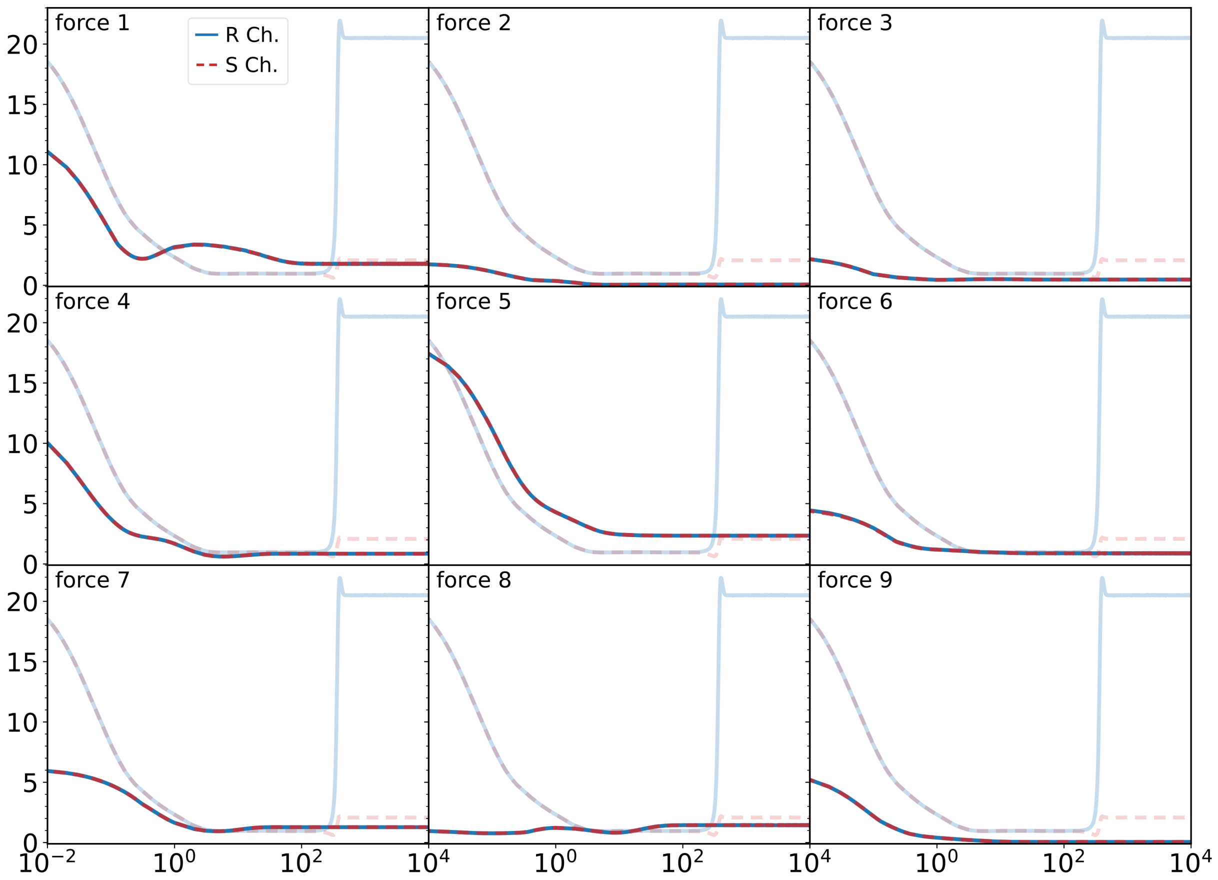


FIG. 12. Work rate plots as a function of time for 9 different randomly drawn forcings of the original model in figure 2b showing that these generally result in racemic, weakly-driven steady-states. Blue solid line indicates $R$ channel while red dotted line indicates the $S$ channel. Background curves in lighter shade illustrate work rate plots for the model with the original forcing which does promote chiral symmetry breaking.

On the other hand, the stochiometric definition in the second work is much broader and allowed the authors to characterize minimal motifs that map to "net" reactions whose products has the overall effect of enhancing the reaction. Such analysis is rooted on a subset decomoposition of the stoichiometric matrices and is therefore dependent on the inherent wiring of the reaction network. However, as we demonstrate next, induction in our model is not a stoichiometric property but is instead dependent on the highly non-linear, and dynamic nature of the driving. As a result, while autocatalysis and autoinduction represent clear manifestations of positive feedback in a chemical system, we maintain our model is best seen through the wider lense of autoinduction.

Importantly, we emphasize that this mode of feedback in our model is the result of enviromental reinforcement through the applied forces. Otherwise, chiral symmetry breaking would manifest through a pre-determined wiring of the system like e.g. autocatalytic domains, cycles etc and without much influence from the drive beyond maintaining a basic energy difference, like that in the Frank model. In fact, we can readily demonstrate this is indeed not the case by swapping the original forces of a chiral symmetry breaking system and re-simulating with another set of randomly-generated forces driving the same set of reactions. We do this for the representative non-racemic system of figure 2b and re-ran against nine other, randomly-drawn forces, as shown in figure 12 for the resulting work rates. Thus, while the original forces induce chiral symmetry breaking, for other types of driving the system not only conserves symmetry but yields unremarkable, weakly-driven steady-states. Similar results can be shown for other models. This test then clearly demonstrates that positive reinforcement, resulting in far-from-equilibrium steady states, must arise from strongly-matched state dependent driving and not from a pre-configured reaction stoichiometry. Without the enviromental feedback of the model, far-from equilibrium states let alone those breaking chiral symmetry, would simply not arise.

Given these considerations, for what concerns the results of our work we thefore conclude that 1) the feedback emerges and is sustained through external forcings, not through an inherent stoichiometric property of the system; 2) it is a state-dependent, non-linear process hence explicitly conditioned on dynamical history with the drives; and remark that 3) the mechanistic details of such forcing are deliberately abstract to encompass a wide-variety of possible scenarios beyond exact chemical properties, and need only reflect energy input changes. For instance, consider a light driven reaction whose products help dissolve suspended, scattering particles and hence increase energy input. In such a case, no complex chemical changes in the reaction (transitions states, intermediates etc) were required, yet it is capable of inducing its own production. For these reasons, we maintain our model should considered in the general perspective of enviromental feedback, and beyond the more limited window of chemical kinetics. In this manner, the exact chemical characterization of a system is not as critical, so long as it is subject to a complex enviromental driving which, in some special circumstances, might yield emergent states strongly conditioned on the exploitation of the energy source. That in some cases such process may also induce spontaneous chiral symmetry breaking is thus indeed one of the main messages of this work.
